# Structure-activity studies of *Streptococcus pyogenes* enzyme SpyCEP reveal high affinity for CXCL8 in the SpyCEP C-terminal

**DOI:** 10.1101/2021.11.14.468564

**Authors:** Max Pearson, Carl Haslam, Andrew Fosberry, Emma J Jones, Mark Reglinski, Robert J. Edwards, Richard Ashley Lawrenson, Jonathan C Brown, Danuta Mossakowska, James Edward Pease, Shiranee Sriskandan

## Abstract

The *Streptococcus pyogenes* cell envelope protease (SpyCEP) is vital to streptococcal pathogenesis and disease progression. Despite its strong association with invasive disease, little is known about enzymatic function beyond the ELR^+^ CXC chemokine substrate range. As a serine protease, SpyCEP has a catalytic triad consisting of aspartate (D151), histidine (H279), and serine (S617) residues which are all thought to be mandatory for full activity. We utilised a range of SpyCEP constructs to investigate the protein domains and catalytic residues necessary for enzyme function. We designed a high-throughput mass spectrometry assay to measure CXCL8 cleavage and applied this for the first time to study the enzyme kinetics of SpyCEP.

Results revealed a remarkably low Michaelis-Menton constant (K_M_) of 82 nM and a turnover of 1.65 molecules per second. We found that an N-terminally-truncated SpyCEP C-terminal construct containing just the catalytic dyad of H279 and S617 was capable of cleaving CXCL8 with a similar K_M_ of 55 nM, albeit with a reduced substrate turnover of 2.7 molecules per hour, representing a 2,200- fold reduction in activity. We conclude that the SpyCEP C-terminus plays a key role in high affinity substrate recognition and binding, but that the N-terminus is required for full catalytic activity.

## Introduction

Group A Streptococcus (GAS) or *Streptococcus pyogenes* is a leading human pathogen that manifests clinically as a broad spectrum of diseases, ranging from less severe, usually self-limiting infections to life-threatening invasive diseases such as necrotizing fasciitis and toxic shock syndrome. While much of the global health burden of *S. pyogenes* can be attributed to auto-immune sequelae such as rheumatic heart disease, invasive diseases contribute greatly to *S. pyogenes*-associated global mortality. At least 663,000 new cases of invasive infection arise per year, accounting for approximately 163,000 deaths [1] with mortality rates remaining high despite intervention; indeed, 20% of patients with invasive *S. pyogenes* disease die within 7 days of infection onset [2].

Several virulence factors contribute to pathogenesis in invasive *S. pyogenes* disease, chief among which is *<u>S</u>treptococcus <u>py</u>ogenes* <u>c</u>ell <u>e</u>nvelope <u>p</u>rotease (SpyCEP), an immune-modulatory cell wall-associated protease. SpyCEP is responsible for the rapid and efficient cleavage of a distinct group of CXC chemokines comprising CXCL1, CXCL2, CXCL3, CXCL5, CXCL6, CXCL7, and CXCL8, both locally at the site of an infection, and systemically [3-6]. The ELR^+^ chemokines, named for their conserved N-terminal glutamate-leucine-arginine motifs, specifically act upon neutrophils eliciting their recruitment and activation. SpyCEP inactivates these chemokines by cleaving the chemokine C-terminal α-helix, releasing a 13 amino acid peptide in the case of CXCL8 [3]. Although chemokine specificity for binding neutrophil chemokine receptors, CXCR1 and CXCR2, is conferred to a large extent by the chemokine N-terminal ELR motif [7], the chemokine C-terminus is necessary for efficient receptor binding and activation [8], in addition to chemokine translocation from tissues to endothelial lumen [7]. As such, SpyCEP cleavage of CXC chemokines results in a reduction of CXCR1 and CXCR2-mediated neutrophil chemotaxis and subsequent paucity of neutrophils at the site of *S. pyogenes* infection [3]. SpyCEP has recently been shown to cleave the human anti-microbial peptide, LL-37 in addition to chemokines [9]. Whilst cleavage does not affect the antimicrobial action of the peptide, it was reported to reduce LL-37 specific neutrophil chemotaxis [9].

SpyCEP is expressed by *S. pyogenes* as a 1647 amino acid, 180 kDa subtilisin-like serine protease, the crystal structure of which was recently solved to 2.8 Å resolution [10] and further refined to 2.2 Å resolution [11]. It is a member of the S8 subtilase family, members of which are characterised by a catalytic triad consisting of an aspartate, histidine, and serine residue, each surrounded by a region of highly conserved amino acids [12]. SpyCEP is unique among streptococcal proteases in that, during maturation, it is autocatalytically cleaved between residues Q244 and S245 into 2 distinct polypeptides, a 30 kDa N-terminal polypeptide and a 150 kDa C-terminal polypeptide. The two polypeptides harbour the separate residues required for the formation of the catalytic site [6]; the N-terminal fragment contains the catalytic D151 residue, and the C-terminal fragment contains the catalytic H279 and S617 residues. Upon cleavage, the two polypeptides re-associate non-covalently to reconstitute the active enzyme [6]. The recent crystal structures have shed further light upon the structure of SpyCEP, describing 9 separate domains, the first 5 of which contain the catalytic triad necessary for enzymatic activity [10].

SpyCEP has been included as a target antigen in several recent *S. pyogenes* vaccine designs due to its cell surface expression, highly conserved nature, and central role in *S. pyogenes* pathogenesis. Immunisation with SpyCEP successfully elicits a SpyCEP-specific neutralising antibody response, providing protection against systemic bacterial dissemination and reducing disease severity in *S. pyogenes* intramuscular, skin infection models and non-human primate infection models [13-18]. Data that demonstrate vaccine dependence on enzyme inhibition highlight the importance of understanding the enzymatic activity of SpyCEP and the potential to improve upon vaccine or inhibitor design [13].

Despite this, little is known about the catalytic properties of the enzyme, except preliminary structure-function studies [6, 10, 11, 19]. Published data largely focus on the impact of SpyCEP on streptococcal pathogenesis and disease progression. In this study we generated multiple SpyCEP constructs to confirm domains necessary for catalytic activity. We then used mass spectrometry to determine the enzyme kinetics of two SpyCEP constructs for the natural substrate CXCL8.

## Methods

### Cloning and purification of recombinant SpyCEP constructs in *Escherichia coli*

Codon-optimised SpyCEP gene constructs were expressed in *Escherichia coli* using synthetic gene sequences (GenScript) from Spy_0416 in the SF370 *S. pyogenes* M1 genome [6, 20] representing full-length enzyme, SpyCEP^34-1613^, the N- terminal polypeptide, SpyCEP^34-244^, and C-terminal polypeptide SpyCEP^245-1613^. Constructs were also generated with alanine substitutions to replace catalytic residues in the N-terminal fragment (SpyCEP^34-244^ D151A), and the C-terminal fragment (SpyCEP^245-1613^ S617A). To enable downstream protein purification, both the full-length enzyme and C-terminal polypeptides were expressed with a C-terminal 6-histidine tag and the N-terminal polypeptides were expressed with an N-terminal FLAG tag and TEV linker.

All SpyCEP constructs were cloned into the vector pET-24B and expressed in BL21 (DE3) competent *E. coli*, cultured in Terrific Broth medium supplemented with 50 µg/mL kanamycin at 37°C and shaken at 200 rpm, for 3 hours. The cultures were induced with 0.5 mM Isopropyl β-D-1- thiogalactopyranoside (IPTG), cooled to 15°C and shaken at 200 rpm for 16 hours before lysis by sonication on ice. Full-length and C-terminal constructs were purified by Ni-IDA affinity chromatography (GenScript) as per the manufacturer’s instructions. N-terminal constructs were purified by anti-flag M2 agarose resin chromatography (Sigma-Aldrich) as per manufacturer’s instruction, and further concentrated and purified by SP ion-exchange (Sigma-Aldrich) and Q FF ion-exchange chromatography (GE Healthcare) as per manufacturer’s instruction. All SpyCEP constructs were subsequently concentrated and purified by size exclusion chromatography on a HiLoad Superdex 200 or 75 prep grade column (GE Healthcare) dependent on molecular weight.

The resultant panel of recombinantly expressed SpyCEP constructs is shown in Table 1. All of the resultant protein preparations were found to be >80% pure as determined following analysis by SDS-PAGE. The re-association of the SpyCEP N and C-termini to create full-length constructs was carried out by equimolar co-incubation at 37°C for 30 minutes in 40 mM Tris-HCl pH 7.5, 0.1 mM CHAPS, 1 mM DTT, 0.1% BSA, 75 mM NaCl.

**Table 1.** SpyCEP constructs used in this study. Constructs were expressed recombinantly in *E. coli* except for s.pSpyCEP that was expressed in, and purified from *S. pyogenes*.

| SpyCEP construct | SpyCEP construct description |
| --- | --- |
| **SpyCEP** | Full-length, SpyCEP<sup>34-1613</sup>-6His |
| **SpyCEP<sup>D151A, S617A</sup>** | Full-length double mutant, SpyCEP<sup>34-1613 D151A, S617A</sup>-6His |
| **N-terminal** | N-terminal fragment, FLAG-TEV-SpyCEP<sup>34-244</sup> |
| **C-terminal** | C-terminal fragment, SpyCEP<sup>245-1613</sup>-6His |
| **N-terminal<sup>D151A</sup>** | N-terminal mutant fragment, FLAG-TEV-SpyCEP<sup>34-244 D151A</sup> |
| **C-terminal<sup>S617A</sup>** | C-terminal mutant fragment, SpyCEP<sup>245-1613 S617A</sup>-6His |
| **s.pSpyCEP** | SpyCEP from *S. pyogenes* supernatant, SpyCEP<sup>34-1613</sup>-6His |

### Expression of histidine-tagged SpyCEP in *S. pyogenes*

To permit release of soluble His-tagged SpyCEP by *S. pyogenes*, the C-terminal anchor domain of SpyCEP was replaced by a stop codon in *S. pyogenes* strain H292, an *emm*81 strain that makes abundant SpyCEP [5]. A 549 base pair region immediately upstream of the SpyCEP cell wall anchor motif was amplified from *S. pyogenes* H292 genomic DNA using the primers 5ʹGGGAATTCTGTTGTCAGGTAACAGTCTTATCTTGCC -3ʹ and 5ʹCCGAATTCACAACACTAGGCTTTTGCTGAGGTCGTTG -3ʹ. *EcoRI* restriction sites were incorporated at the terminal ends of the primer sequences. The amplified DNA was cloned into the homologous recombination plasmid pUCMUT to produce the vector pUCMUT_CEP_ which was transformed into One Shot TOP10 Chemically Competent *E. coli* (Thermo Fisher Scientific) according to the manufacturer’s instructions. The nucleotide sequence of the SpyCEP C-terminus was subsequently further modified to encode a hexa-histidine tag by inverse PCR using pUCMUT_CEP_ as the template and the primers 5’- TATCCTAGGTAGTGTTGTGAATTCGTAATCATGGTCATAG-3’ and 5’- TATCCTAGGATGATGATGATGATGATGGGCTTTTGCTGAGGTCGTTG-3’. The amplification was performed using GoTaq Long PCR Master Mix (Promega) according to the manufacturer’s instructions. *AvrII* restriction sites, incorporated at the terminal ends of the primer sequences, facilitated re-circulation of the amplified plasmid. The presence of a 6-His sequence in the modified pUCMUT_CEP_ construct (denoted pUCMUT_CEP-HIS_) was confirmed by Sanger sequencing using the pUCMUT sequencing primers 5’-GACAGCAACATCTTTGTGAAAGATGG-3’ and 5’- CATTAATGCAGCTGGCACGAC-3’. The pUCMUT_CEP-HIS_ construct was introduced into H292 by electroporation and crossed into the chromosome by homologous recombination as previously described [21] to generate strain H1317. Secretion of 6-His-tagged SpyCEP into the culture supernatant of H1317 was confirmed by western blotting (data not shown). To purify SpyCEP, *S. pyogenes* H1317 was grown in Todd-Hewitt broth (Oxoid) overnight at 37°C with 5% C0_2_ The culture was pelleted at 2500 xg for 10 minutes and the supernatant sterilised using Amicon 0.22 µm filters, then concentrated using 15 ml Amicon 100 kDa centrifuge filter columns and purified by nickel column affinity chromatography (Novagen His-Bind Resin) as per the manufacturer’s instructions.

### Sodium dodecyl sulphate–polyacrylamide gel electrophoresis (SDS-page)

To visualise chemokine cleavage, 50 pmol of recombinant human CXCL1 or CXCL8 (R&D Systems) was incubated with C-terminal SpyCEP in a final volume of 20 μl, at a molar ratio of 1:5, 50, 500, 5000 in favour of chemokine for 2 hours at 37^°^C. C-terminal^S617A^ and N-terminal constructs^34-244^ were included as controls and assayed at the highest 1:5 molar ratio. Reactions were halted by the addition of Dithiothreitol (DTT) at a final concentration of 100 mM, 4X Bolt LDS sample buffer and heating to 70°C for 10 minutes. Samples were separated on pre-cast 4-12% MES buffered Bolt Bis-Tris gradient gels (Invitrogen) by SDS-PAGE gel electrophoresis at 165 V for 35 minutes with SeeBlue Plus 2 (Invitrogen) used for molecular weight ladder. Gels were stained with PageBlue protein staining solution (Thermo Fisher Scientific) overnight and de-stained in deionised water.

### Multiplex fluorescent western blotting

To visualise generation of both intact CXCL8 and the larger (N-terminal) CXCL8 cleavage product, a rabbit antiserum was raised against the CXCL8 neo-epitope (anti-ENWVQ) that is exposed following SpyCEP cleavage of CXCL8 [22]. SpyCEP constructs were incubated at 37 °C with 18.75 pmol of recombinant human CXCL8 in a final volume of 20 μl, at a 1:50 molar ratio in favour of CXCL8. Digests were halted and separated by SDS-PAGE gel electrophoresis as described above. Proteins were transferred by iBlot2 (Thermo Fisher Scientific) onto 0.2 μm nitrocellulose membranes as per manufacturer’s instructions. Membranes were subsequently blocked for 1 hour at room temperature in blocking buffer (PBS with 5% skimmed milk powder (Sigma-Aldrich) and 0.1% Tween), then blotted overnight at 4 °C with 2 primary antibodies, 1 µg/ml mouse anti-human CXCL8 (R&D Systems) and 1:1000 rabbit anti-ENWVQ. Membranes were washed in wash buffer (PBS with 0.05% Tween) and incubated with 1:7500 goat anti-rabbit IgG A680nm and 1:7500 goat anti-mouse IgG A790nm for 1 hour at room temperature before being visualised on LiCor Odyssey Fc (Invitrogen).

SpyCEP activity against LL-37 was assessed by a 16 hour, 37 °C incubation of 111.1 pmol human LL-37 (R&D Systems) with SpyCEP constructs at a 1:10 molar ratio in favour of LL-37 in a final volume of 20 μl. Reactions were stopped, separated and blotted onto 0.2 μm nitrocellulose as above and incubated overnight at 4°C in blocking buffer supplemented with 2 μg/ml polyclonal sheep IgG anti-LL-37 (R&D Systems). Membranes were washed in wash buffer (PBS with 0.05% Tween) and incubated in blocking solution with rabbit anti-sheep IgG (Abcam) at 1:40,000 dilution for 1 hour at room temperature and visualised on LiCor Odyssey Fc (Invitrogen).

### Enzyme linked immunosorbent assay (ELISA) analysis of chemokine cleavage

Catalytic activities of SpyCEP constructs were measured through detection of remaining intact CXCL8 or CXCL1 substrate following incubation with SpyCEP by ELISA (R&D Systems human CXCL8 and CXCL1 DuoSet ELISA) as per manufacturer’s instructions. For cleavage time courses of CXCL8 by C-terminal SpyCEP, 5 pmol CXCL8 were incubated with C-terminal constructs at a range of enzyme: chemokine molar ratios, 1:5 – 1:250, in a final volume of 100 μl at room temperature for 60 minutes. Full-length SpyCEP and C-terminal ^S617A^ were included as controls at a molar ratio of 1:1000 and 1:5, respectively. To compare SpyCEP cleavage rates of CXCL8 and CXCL1, 5 fmol or 10 fmol SpyCEP were incubated with 2 pmol human CXCL8 and CXCL1 respectively (R&D Systems), in a final volume of 100 μl and incubated at room temperature for 30 minutes. All reactions were halted at defined timepoints with the addition of concentration of Pefabloc (Sigma-Aldrich) to a final concentration of 2 mg/ml (8.34 mM).

Linear regression analyses of the initial five time points of CXCL1 and CXCL8 cleavage (0, 1, 2, 3 and 4 minutes) were utilised to determine the maximal rate of SpyCEP activity.

### Mass spectrometry analysis of CXCL8 cleavage

Analysis of CXCL8 cleavage was assayed on a SCIEX API6500 triple quadrupole electrospray mass spectrometer coupled to a high-throughput robotic sample preparation and injection system, RapidFire200 (Agilent Technologies). CXCL8 substrate, the 13 amino acid CXCL8 SpyCEP cleavage product RVVEKFLKRAENS, and a heavy atom substituted internal standard of CXCL8 SpyCEP cleavage product RV [U_13_C_5 15_N-VAL]-EKF-[U-_13_C_6 15_N-Leu]- KRAENS) were monitored by mass spectrometry. The mass spectrometer was operated in positive electrospray MRM mode, and transitions (Q1/Q3) for each species were optimised to give m/z as follows: CXCL8, 1048.7/615.3, CXCL8 cleavage product 526.2/211.1, internal standard 530.5/211.1. A dwell time of 50 ms was used for the MRM transitions. The mass spectrometer was operated with a spray voltage of 5500 V and at a source temperature of 650 °C.

To assay CXCL8 cleavage dynamically, chemokine and SpyCEP constructs were loaded into a 384 well plate to the following final concentrations: 6.25-2000 nM chemokine, 250 pM SpyCEP or 40 nM C-terminal SpyCEP in a 20 μl reaction volume. Reactions were carried out at 21°C and stopped at desired time points, 0-240 minutes, by the addition of stop solution (1% formic acid) supplemented with 1 µM heavy atom substituted CXCL8 internal standard and centrifuged 2000 xg for 10 minutes. Assay plates were transferred onto the RapidFire200 integrated to the API6500 mass spectrometer. Samples were aspirated under vacuum directly from 384-well assay plates for 0.6 s. The samples were then loaded onto a C18 solid-phase extraction cartridge to remove non-volatile buffer salts, using HPLC -grade water supplemented with 0.1% (v/v) formic acid at a flow rate of 1.5 mL/min for 4 s. The retained analytes were eluted to the mass spectrometer by washing the cartridge with acetonitrile HPLC-grade water (8:2, v/v) with 0.1% (v/v) formic acid at 1.25 mL/min for 4 s. The cartridge was re-equilibrated with HPLC-grade water supplemented with 0.1% (v/v) formic acid for 0.6 s at 1.5 mL/min.

Results were expressed as a ratio of CXCL8 cleavage product (526.2/211.1) intensity areas compared with the internal standard (530.5/211.1) instensity areas. The results were interpolated from a standard curve constructed using known amounts of cleaved CXCL8 over the range of 25 – 2000 nM where there was a linear relationship (R^2^=0.99) and with a coefficient of variation < 9% for all concentrations. Where interpolated values were below 25 nM, the assay LOQ, values were assigned an arbitrary value of LOQ/2, 12.5 nM.

### Kinetic analysis

Linear regression analyses from five time points of full-length SpyCEP (1, 2, 3, 4, 5 minutes) and C-terminal SpyCEP (15, 30, 45, 60, 75 minutes) CXCL8 reactions were plotted against substrate concentration and kinetics derived from the Michaelis-Menton equation Y = V_max_*X/ (K_M_ + X) and the K_cat_ equation Y = ET*k_cat_*X/(K_M_ + X) where Et = enzyme concentration as fitted by GraphPad Prism v 8.0.2.

## Results

### Screening of CXCL8 cleavage activity using fluorescent western blotting

To initially assess the activity of recombinant SpyCEP constructs we screened constructs for specific CXCL8 cleavage activity using two-colour multiplex western blotting. Human CXCL8 was incubated for 2 hours at 37 °C with the different SpyCEP constructs, and the reaction products were separated by SDS-PAGE, then immunoblotted using separate antibodies that detect either intact or cleaved CXCL8. Detection of full-length CXCL8 (8 kDa, green bands) or the CXCL8 neo epitope (ENWVQ), exposed after SpyCEP cleavage, (6 kDa, red bands) was evident with this system (Figure 1). Recombinant full-length SpyCEP expressed in *E. coli* successfully cleaved CXCL8 to completion, with no difference observed between CXCL8 cleavage with *S. pyogenes* SpyCEP and *E. coli* SpyCEP (Figure 1). As expected, no CXCL8 cleavage was observed when using the catalytically inactive mutant, SpyCEP^D151A, S617A^. As has been previously reported [6], the SpyCEP N- and C-termini, when independently expressed and purified, can be re-associated to form an active enzyme R.E SpyCEP, which successfully cleaved CXCL8. The N-terminal fragment of SpyCEP alone could not cleave CXCL8. Unexpectedly, however, the C-terminal fragment of SpyCEP was observed to cleave CXCL8, albeit not to completion. This independent catalytic activity was negated by mutation of the catalytic S617 to alanine (C-terminal^S617A^ construct, Figure 1). Cleavage of CXCL8 by the SpyCEP C-terminal fragment was enhanced when re-associated with the catalytically inert N-terminal mutant (N-terminal^D151A^ construct); indeed, activity was equivalent to that observed with the re-associated enzyme under these conditions, with cleavage of CXCL8 to near completion. However, the N-terminal^D151A^ construct was unable to restore catalytic activity to the C-terminal^S617A^ construct when the two were combined.

**Figure 1.**
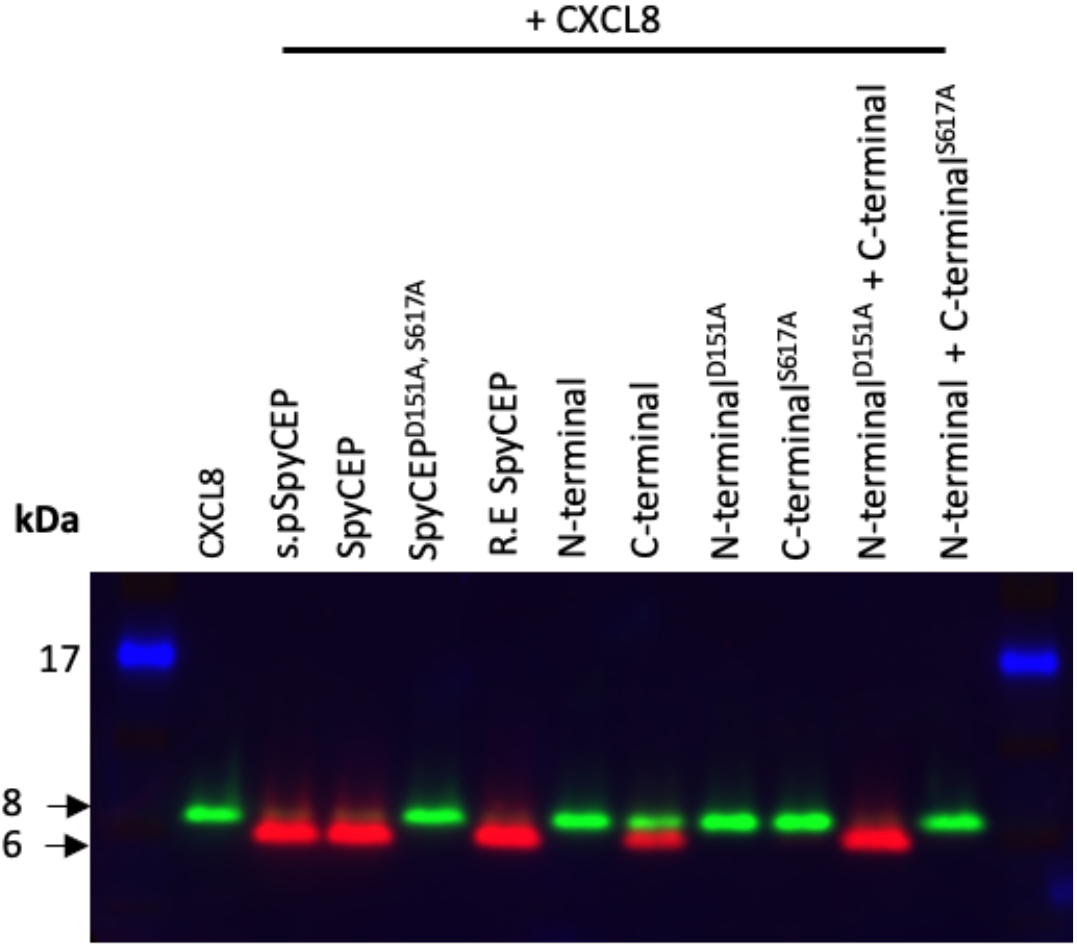
Cleavage activity of recombinant SpyCEP constructs assayed by immunoblot. Two colour immunoblot showing cleavage of 18.75 pmol CXCL8 when incubated for 2 hours at 37 °C either alone (2^nd^ lane) or with a panel of SpyCEP constructs at a 1:50 molar ratio (SpyCEP: CXCL8). Green bands represent intact CXCL8 (anti-CXCL8 antibody); red bands represent cleaved CXCL8 (anti-ENWVQ). A 17 kDa molecular weight marker is shown in blue, 8 kDa and 6 kDa molecular weights are highlighted by arrows. Figure is representative of 2 independent immunoblots.

The results demonstrated for the first time that the SpyCEP C-terminal fragment alone is sufficient for catalytic cleavage of CXCL8 in this assay system. The data also established that, although the presence of the SpyCEP N-terminal fragment enhanced enzymatic activity, the N-terminal residue D151 was dispensable for enzymatic activity. We further sought to examine whether the C-terminal displayed activity against the newly described substrate, LL-37 [9]. Western blot analysis confirmed cleavage of LL-37 by full-length recombinant SpyCEP, demonstrated by a reduction in band size, however the C-terminal fragment was unable to cleave LL-37 despite a high molar ratio of enzyme to LL-37 (1: 10) and a prolonged 16-hour incubation at 37 °C (Figure S1).

### Differential cleavage of CXCL8 and CXCL1 by full-length and C-terminal SpyCEP

To determine whether the catalytic activity of the C-terminal SpyCEP fragment was reproducible over a shorter incubation period, we assessed CXCL8 cleavage by ELISA, where CXCL8 cleavage is detected through a reduction in substrate concentration. SpyCEP C-terminal constructs were incubated with CXCL8 at molar ratios ranging from 1:5 – 1:250 (enzyme: chemokine) over a 60- minute time course at room temperature. Full-length recombinant SpyCEP and the inactive C-terminal fragment, C-terminal^S617A^, were included as controls at a molar ratio of 1:1000 and 1:5, respectively. Under these conditions, near complete CXCL8 cleavage was observed for full-length SpyCEP and a dose-dependent increase in catalytic activity was observed for the C-terminal fragment (Figure 2). After 5 minutes incubation full-length SpyCEP cleaved over 50% of the starting CXCL8 input, with only 6.25% of CXCL8 remaining after 1 hour. At the highest concentration of C-terminal SpyCEP tested, a 1:5 molar ratio, the C-terminal fragment alone cleaved 9% of the starting CXCL8 by 5 minutes, 25% by 30 minutes and 42% by 60 minutes. Indeed, at the lowest concentration tested, a 1:250 molar ratio, the C-terminal of SpyCEP cleaved 9% of CXCL8 input by 1 hour. As demonstrated by immunofluorescent western blotting, the serine residue at position 617 was vital for SpyCEP catalytic function, as no CXCL8 cleavage was observed using the C-terminal^S617A^ construct, even when employed at the highest enzyme: chemokine ratio. After a 1-hour incubation, full-length SpyCEP and C-terminal constructs (assayed using a molar enzyme: chemokine ratio of 1:5, 1:25 and 1:50) cleaved significantly more CXCL8 compared to the C-terminal^S617A^ construct.

**Figure 2.**
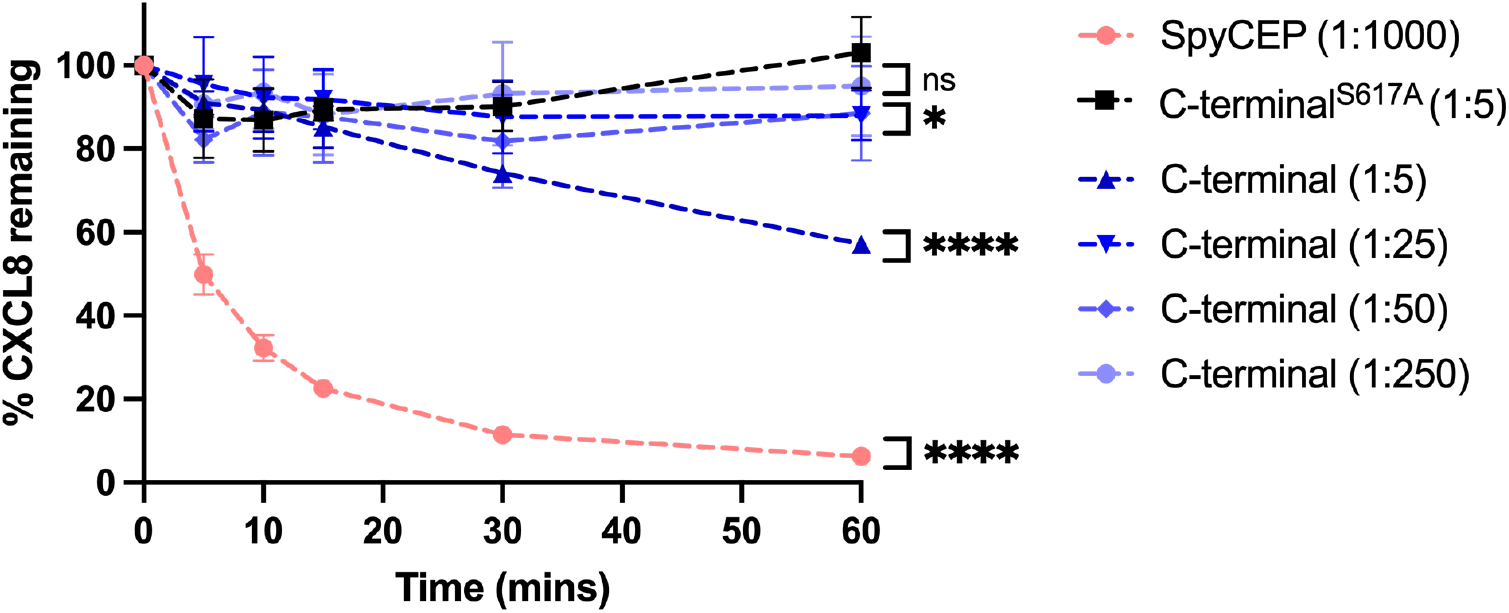
Cleavage activity of SpyCEP and C-terminal SpyCEP^245-1613^ constructs using CXCL8 ELISA. SpyCEP constructs were co-incubated with 5 pmol CXCL8. Graphs show residual CXCL8 after a 60- minute room temperature incubation, using full-length SpyCEP at a 1:1000 ratio to CXCL8; the C-terminal SpyCEP construct at a 1:5 – 1:250 molar ratio to CXCL8; and the C-terminal^S617A^ mutant at a 1:5 molar ratio to CXCL8. Reactions were halted at specified timepoints by the addition of Pefabloc to a final concentration of 2 mg/ml. N=6 experimental replicates for each construct, data points show means, error bars represent SD. ns p > 0.05, * p ≤ 0.05, **** p ≤ 0.0001, at 60 minutes as determined by ordinary one-way ANOVA.

SDS-PAGE analysis of the catalytic activity of the C-terminal SpyCEP construct additionally showed that C-terminal activity was not restricted to the CXCL8 substrate. Over 2 hours at 37 °C using a 1:5 molar ratio of enzyme: substrate, the SpyCEP C-terminal construct was capable of cleaving human CXCL1 to near completion (Figure S2).

To further assess the activity of SpyCEP and to interrogate reaction rates against separate chemokines, full-length SpyCEP was incubated with CXCL1 and CXCL8 and the remaining chemokine levels determined by ELISA. Human CXCL1 or human CXCL8 were incubated with SpyCEP at room temperature over a 30-minute time course, at 1:200 or 1:400 molar ratios respectively (enzyme: chemokine). 5 fmol of SpyCEP rapidly and efficiently cleaved 2 pmol CXCL8, with ∼ 15% of the chemokine input remaining after 10 minutes of incubation (Figure 3A). This contrasted with CXCL1 cleavage, that required 10 fmol SpyCEP to cleave just 25% of the chemokine input over the same 10- minute period. Indeed, by 30 minutes SpyCEP had yet to cleave half of the starting amount of CXCL1 (Figure 3B). Utilising a linear regression of the initial 5 timepoints, where SpyCEP activity is maximal, we found that 5 fmol of SpyCEP was able to cleave 284 fmol of CXCL8 per minute, and 10 fmol of SpyCEP was capable of cleaving 62 fmol of CXCL1 per minute. These data confirmed the activity of recombinant SpyCEP and highlighted differential cleavage efficiency across the CXC substrate range – a feature which has been previously recognised but not quantified [8].

**Figure 3.**
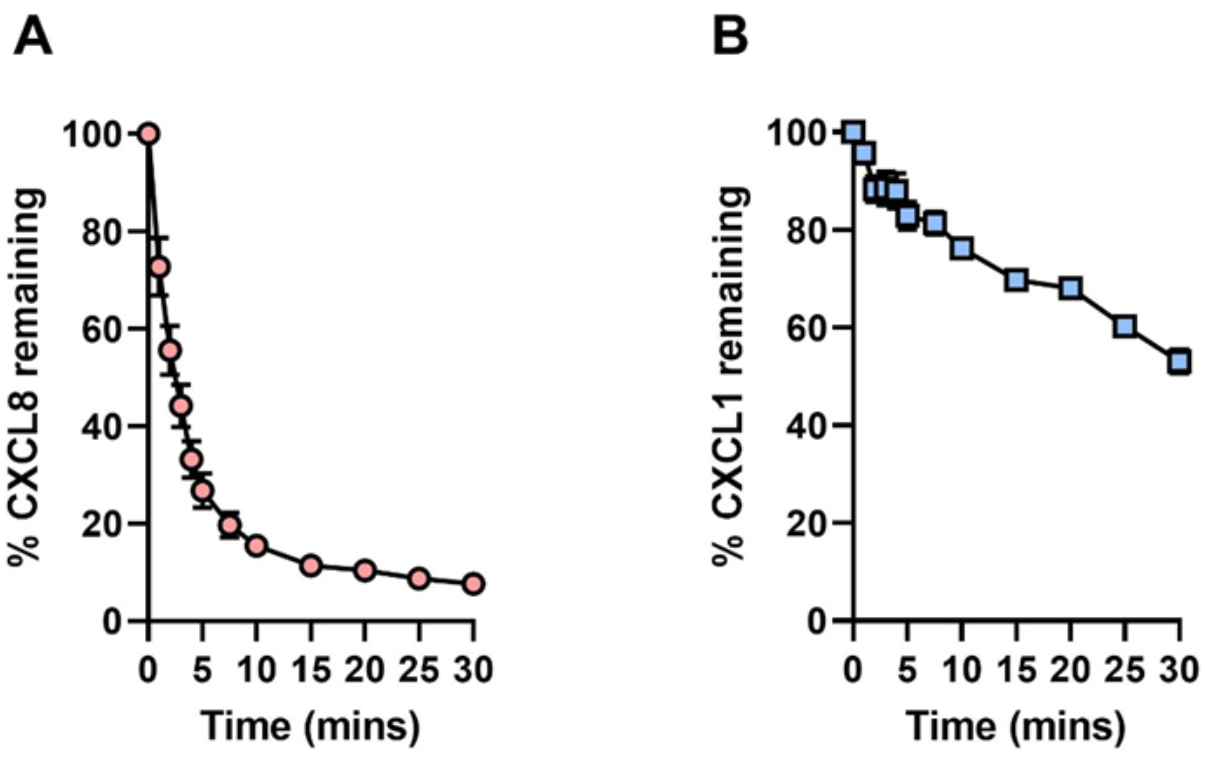
Cleavage of CXCL8 and CXCL1 by recombinant full-length SpyCEP. 30-minute, room temperature cleavage time course of: A. 2 pmol CXCL8 with 5 fmol SpyCEP and B. 2 pmol CXCL1 with 10 fmol SpyCEP. Cleavage reactions were halted at specified timepoints by the addition of Pefabloc to a final concentration of 2 mg/ml and the remaining chemokine measured by ELISA. N=6 experimental replicates for each data point, error bars represent SD of mean values.

### Mass spectrometry-derived kinetics of active SpyCEP constructs

To directly compare the active full-length and C-terminal SpyCEP constructs and to understand relative catalytic efficiencies, a mass spectrometry approach to continuously assay the generation of the 13 amino acid peptide cleaved from the native substrate CXCL8, following incubation with enzyme, was employed. A range of CXCL8 concentrations (25-2000 nM) were incubated with a fixed concentration of enzyme, either 250 pM of full length SpyCEP, or 40 nM of the C-terminal SpyCEP^245-1613^ and the production of the 13 amino acid peptide was monitored over time. 250 pM full-length SpyCEP cleaved CXCL8 to near completion over 30 minutes when substrate concentrations were less than 250 nM; incomplete CXCL8 cleavage was observed when substrate concentration was in excess of 250 nM (Figure S3A). In contrast, 40 nM of the C-terminal SpyCEP^245-1613^ cleaved CXCL8 to near completion over 4 hours when the CXCL8 concentration was 250 nM or less; incomplete CXCL8 cleavage was again observed when substrate concentrations were over 250 nM (Figure S3B). Under these conditions the C-terminal SpyCEP fragment maintained measurable catalytic activity throughout, though with reduced efficacy compared to the full-length construct. A 160-fold increase in enzyme concentration and additional 3.5 hours reaction time were required to cleave a comparable amount of CXCL8.

Linear regression analyses of 5 time points, where the rate of cleaved CXCL8 production was linear, were used to derive Michaelis-Menton plots (Figure 4) and K_M_ and K_cat_ values for each construct (Table 2).

**Figure 4.**
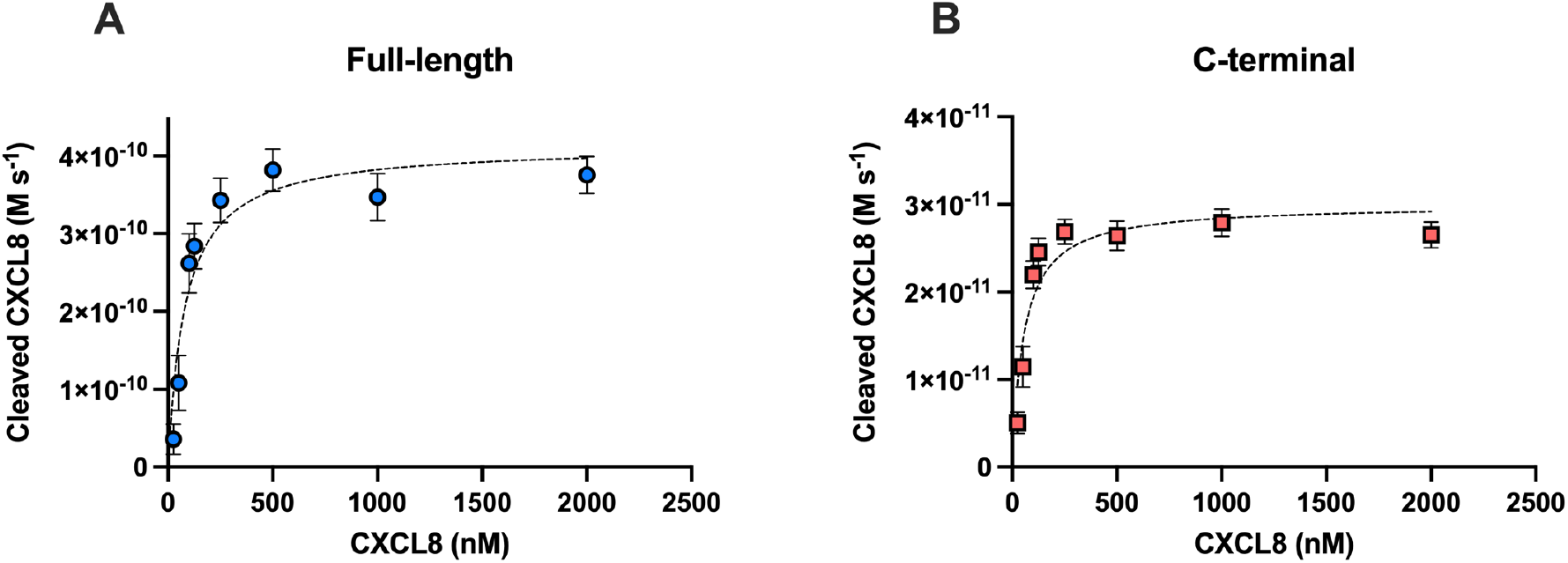
Kinetic activity of full-length and C-terminal SpyCEP construct cleavage of CXCL8. Data points represent the mean change in cleaved CXCL8 production over time (M s^-1^) of, **A.** Full-length SpyCEP and **B.** C-terminal SpyCEP. Error bars represent the standard error of the mean of 5 reactions.

**Table 2.** Kinetic parameters of full-length and C-terminal SpyCEP construct activity in cleavage of CXCL8. K_cat_, K_M_ and V_max_ ± SEM and K_cat_ / K_M_ for full-length SpyCEP and C-terminal SpyCEP derived from Michaelis-Menton graphs Y = V_max_*X/ (K_M_ + X) and K_cat_ equation Y = ET*k_cat_*X/(K_M_ + X) where Et = enzyme concentration, 2.5 x10^-10^ M for full-length SpyCEP and 4 x10^-8^ M for C-terminal SpyCEP. K_M_ values for full-length SpyCEP and the C-terminal fragment were 82 nM and 55 nM respectively, suggesting nanomolar affinity of the SpyCEP constructs for CXCL8. The similarity in the K_M_ values between the C- terminal SpyCEP fragment and the full-length enzyme point to a similar ability to bind substrate. The K_cat_ values (indicative of the number of molecules of cleaved CXCL8 produced per second) however, differed substantially. The full-length enzyme cleaved 1.65 molecules of CXCL8 per second compared to the C-terminal construct which cleaved 2.7 molecules per hour, a 2,200-fold reduction in activity.

| Construct | $K_{cat} \pm SEM (s^{-1})$ | $K_M \pm SEM (nM)$ | $V_{max} \pm SEM (M s^{-1})$ | $K_{cat}/K_M (M^{-1} s^{-1})$ |
| --- | --- | --- | --- | --- |
| Full-length | $1.65 \pm 0.13$ | $82 \pm 26$ | $4.14 \times 10^{-10} \pm 3.35 \times 10^{-11}$ | $2.02 \times 10^7$ |
| C-terminal | $0.00075 \pm 0.00005$ | $55 \pm 17$ | $3.00 \times 10^{-11} \pm 2.17 \times 10^{-12}$ | $1.36 \times 10^4$ |

## Discussion

SpyCEP is a serine protease and a leading virulence factor of *S. pyogenes,* with a narrow range of substrate specificity, restricted to the family of ELR^+^ CXC chemokines which modulate neutrophil mediated immune responses [3, 4, 8] and LL-37 [9]. Autocatalytic processing of SpyCEP results in the generation of two individual fragments that re-assemble to form an active enzyme [6]. Here, we describe the enzyme kinetics of full-length SpyCEP and report the K_M_ of the enzyme for its natural substrate to be remarkably low, just 81.76 nM, consistent with high efficiency. Furthermore, we demonstrate that the C-terminal SpyCEP fragment can catalyse the cleavage of CXCL1 and CXCL8 independent of the N-terminal fragment. Indeed, when K_M_ values were compared, they were found to be similar, suggesting that substrate binding may be confined to the C-terminal domain of SpyCEP. The enzymatic activity of the C-terminal SpyCEP fragment was, however, markedly reduced compared to full-length SpyCEP. The N-terminal and N-terminal^D151A^ constructs were equally able to restore full catalytic activity of the C-terminal SpyCEP fragment. Collectively, this suggests that although the aspartate 151 of the N-terminal fragment may be dispensable for catalysis, the domain itself is important for optimal enzyme activity.

Serine proteases are ubiquitous and comprise up to one third of all proteolytic enzymes currently described. They are currently categorised by the MEROPS database [23] into 13 distinct clans, being differentiated into groups of proteins based on their evolution from the same common ancestor, with SpyCEP belonging to the S8 family of the SB clan. S8 serine proteases are typified by a classical catalytic triad composed of serine, histidine and aspartic acid residues that together contribute to the hydrolysis of a peptide bond within the substrate. It is recognised that a number of serine protease clans employ a variation on the S8 catalytic triad, utilising instead a triad of serine, histidine, and glutamic acid, or serine, glutamic acid, and aspartic acid residues. Other clans utilise catalytic dyads of lysine and histidine or histidine and serine for proteolytic activity. Our data suggest that SpyCEP activity can reside in a catalytic dyad of histidine and serine, albeit at a greatly reduced efficacy. It is likely this large gulf in efficiency explains the failure of previous studies to detect catalytic activity within the isolated C-terminal SpyCEP fragment [6].

Within serine proteases there are additional features, beyond the catalytic triad residues, which can contribute to activity. The oxyanion hole for example, a pocket in the active site composed of backbone amide NH groups, may provide substrate stabilisation and help drive catalysis [24]. Additional residues located in close proximity to the catalytic pocket can also mitigate a loss of activity resulting from a missing residue, and water also has the potential to moonlight as a missing functional group [24]. These ‘stand ins’ can provide a possible substitute machinery to help drive catalytic function. Indeed, some studies have shown that, even with all three catalytic triad residues removed, serine proteases are still capable of catalysis at rates 1000-fold greater than the background rate of hydrolysis [24-26].

Mass spectrometry-based kinetics showed that full-length active SpyCEP has a K_M_ of 82 nM and K_cat_ of 1.65 molecules per second; values which are in agreement with our initial ELISA-based kinetic assessment [11]. In contrast, the C-terminal fragment of SpyCEP has a K_M_ of 55 nM and K_cat_ of 0.00075 molecules per second. These constructs both demonstrated low, nanomolar K_M_ values, suggesting a high binding efficiency of SpyCEP for its natural substrate CXCL8, likely conferred by the C-terminal domain. This is in keeping with the fact that low nanomolar concentrations of CXCL8 are optimal for neutrophil recruitment [8]. Although K_M_ values are often reported in the µM – mM range, nanomolar K_M_ values are not without precedent for other serine proteases; human kallikrein 6 has a reported K_M_ of 300 nM and Factor Xa, a constituent of the prothrombinase complex, has a K_M_ of 150 nM for prothrombin [27, 28]. Enzyme specificity, a constant which measures the cleavage efficiency of enzymes, (K_cat_ /K_M_), for full-length SpyCEP was estimated to be 2.02 x10^7^ M^-1^ s^-1^, a value in the order of magnitude typical for serine proteases [24]. The specificity constant of the C-terminal fragment was ∼1500-fold less, 1.36 x10^4^ M^-1^ s^-1^, and K_cat_ 2200 fold less, a reduction that is in line with previously reported aspartic acid mutants from a systematic mutational study of the *Bacillus amyloliquefaciens* subtilisin catalytic triad [26]. The development of an MS-based assay of SpyCEP activity, that is not dependent on western blotting or ELISA, provides potential for future high-throughput analysis of SpyCEP activity and detection of SpyCEP inhibitors.

The kinetic assays attributed a marked increase in CXCL8 turnover to the additional presence of the N-terminal fragment. Although we did not specifically evaluate the role of the aspartate residue at position 151 on the kinetics of SpyCEP activity, this residue did not contribute appreciably to cleavage of CXCL8 when evaluated using immunoblotting. This raises a question as to whether the N-terminal fragment confers some additional structural contribution to enzyme activity. Our data strongly suggest that substrate binding is likely to be attributed to the C-terminal fragment, a finding consistent with related cell envelope proteinases of *Lactococci* [12] and the closely related streptococcal protein, C5a peptidase [29].

The implications of our findings relating to activity of the C-terminal SpyCEP fragment in *S. pyogenes* pathogenesis are currently unclear; without the N-terminus, the enzymatic activity detected may be too low to be of consequence for chemokine cleavage *in vivo,* and in nature both N- and C-terminal fragments are likely to co-exist. SpyCEP has been a focus of *S. pyogenes* vaccine development in recent years [30], used either in isolation or combination with other antigenic targets [11, 14, 16, 17, 31]. Vaccine-induced SpyCEP specific antibodies appear not to act through traditional opsonic means [13] and so may act through inhibition of SpyCEP activity. Many vaccine preparations evaluated have been based on ‘CEP5’; a polypeptide spanning residues 35 – 587 of SpyCEP which contains the N-terminal fragment and only part of the C terminal fragment [18]. Our findings relating to enzyme function suggest that antibodies targeting the C-terminal fragment of SpyCEP are more likely to provide greater neutralizing activity, and potentially improve vaccine efficacy.

## Funding

This work was funded by a Biotechnology and Biological Sciences Research Council (BBSRC) NPIF iCASE studentship and a Wellcome Trust Collaborative award 215539: ‘Understanding and exploiting Group A streptococcal anti-chemotactic proteases in vaccines for infection’.

## Acknowledgements

The authors are grateful for the support of a GSK Golden Triangle Discovery Partnership in Academia Collaboration (DPAC) that permitted development of the mass spectrometry high throughput cleavage assay and reagent provision. SS acknowledges the Imperial College London NIHR Biomedical Research Centre.

## Data availability

No new datasets were generated in this study other than those reported in the manuscript.

## Author contributions

MP, CH, AF, DM, JP and SS designed the study. MP, CH, MR, RJE, RAL and JB collected the data. MP and CH analysed the data. MP, CH, AF, DM, JP and SS interpreted the data. MP collected and prepared the figures. MP, MR, JP and SS drafted the manuscript. MP, CH, AF, MR, RJE, RAL, DM, JP, SS, revised the manuscript content.

## Conflict of interest

Co-authors Carl Haslam, Andrew Fosberry, Emma J Jones and Danuta Mossakowska were employed by GlaxoSmithKline R&D at the time this work was conducted. All other authors declare no conflict of interest.

## Notes

### Summary of Updates

Title change to be more descriptive Abstract re-organised Text amendment in line with reviews. Concentrations and amounts provided in more consistent units. Values of cleaved CXCL8 that were below limit of quantification (LOQ) now assigned a value of LOQ/2, rather than zero, as stated in Methods, and kinetic data recalculated (Table 2).

