## Supplementary figures for "Structure-activity studies of *Streptococcus pyogenes* enzyme SpyCEP reveal high affinity for CXCL8 in the SpyCEP C-terminal"

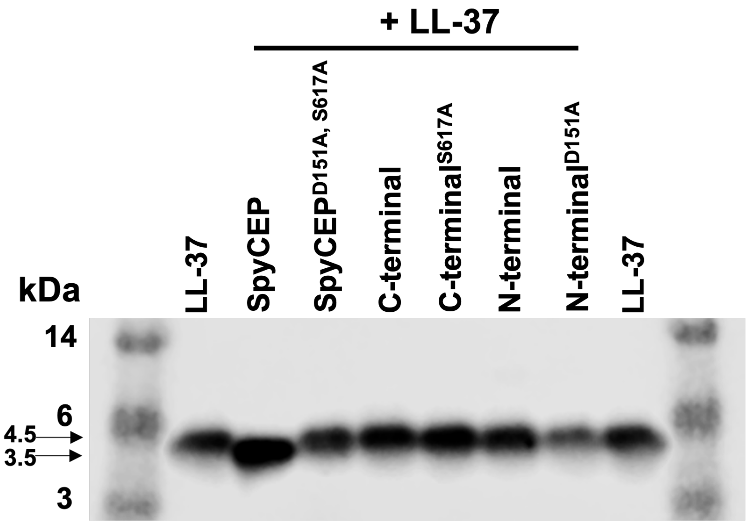


**Figure S1. Western blot of SpyCEP specific LL3-7 cleavage.** Immunoblot of 111.1 pmol human LL-37 incubated for 16 hours at 37 °C either alone (lane 1 and 8) or with a panel of SpyCEP constructs at a 10:1 molar ratio in favour of LL-37. 4.5 kDa full-length LL-37 band and 3.5 kDa cleaved LL-37 band are indicated by arrows and were detected with 2 µg/ml sheep IgG polyclonal LL-37 antibody and rabbit anti-sheep IgG antibody (1:40,000). Figure is representative of two experiments.

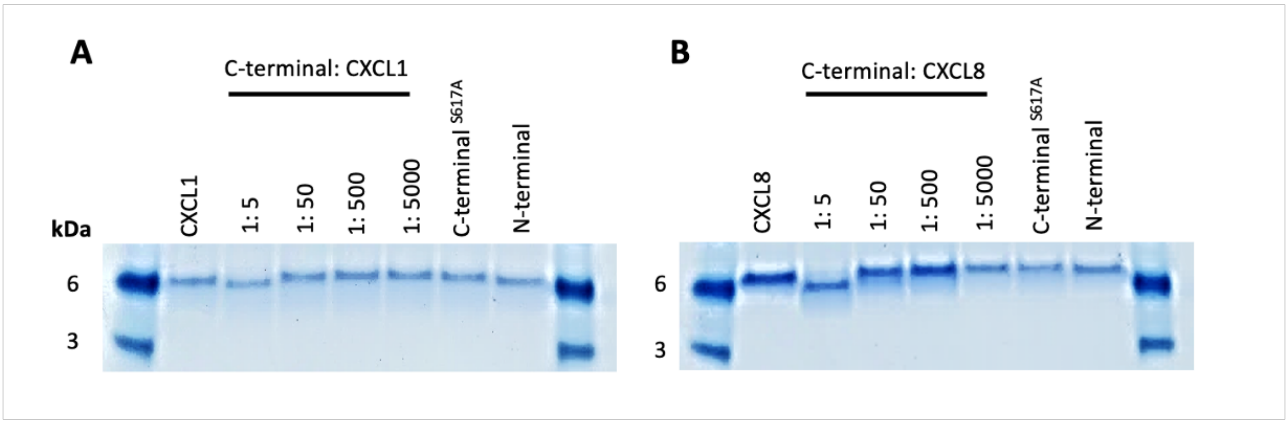


**Figure S2. Comparative cleavage of CXCL1 and CXCL8 by C-terminal SpyCEP.** SDS-page analysis of A. 50 pmol human CXCL1 (all lanes) or B. 50 pmol human CXCL8 (all lanes) cleavage by C-terminal SpyCEP at a 1:5 – 1:5000 molar ratio (SpyCEP: Chemokine). C-terminal$^{S617A}$ and N-terminal controls were all assayed at the highest 1:5 molar ratio.

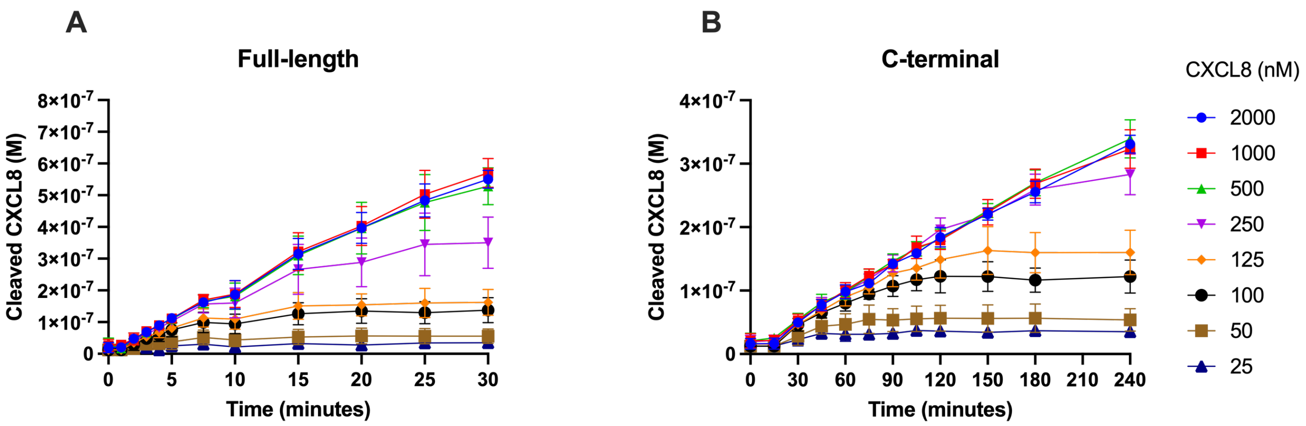


**Figure S3. CXCL8 cleavage by active recombinant SpyCEP constructs using mass spectrometry**. **A.** Production of the 13 amino acid species cleaved from CXCL8 by 250 pM of full-length SpyCEP over 30 minutes at room temperature, following incubation with 6.25 nM – 2000 nM CXCL8, 1: 250 – 8000 molar ratio (SpyCEP: CXCL8). **B**. Production of the 13 amino acid species by 40 nM SpyCEP C-terminal fragment over a 4 hour room temperature incubation, following incubation with 6.25 nM – 2000 nM CXCL8, 1: 0.15 – 50 molar ratio (SpyCEP: CXCL8). N=5 experimental replicates per data point, error bars represent SD. Figure is representative of two experiments
